# Stepwise recombination suppression around the mating-type locus associated with a diploid-like life cycle in *Schizothecium* fungi

**DOI:** 10.1101/2025.04.18.645578

**Authors:** Elsa De Filippo, Elizabeth Chahine, Jeanne Legendre--Despas, Alodie Snirc, Amandine Labat, Pauline Michel, Pierre Grognet, Valérie Gautier, Emilie Levert, Christophe Lalanne, Philippe Silar, Tatiana Giraud, Fanny E. Hartmann

## Abstract

Recombination suppression often evolves around sex-determining loci and extends stepwise, resulting in adjacent regions with different levels of divergence between sex chromosomes, called evolutionary strata. In Ascomycota fungi, evolutionary strata have been found around the mating-type (*MAT*) locus only in pseudo-homothallic species, *i.e*., with a diploid-like lifecycle and mycelia carrying nuclei of both mating types. In contrast, no recombination suppression has been observed in fungi with a haploid-like lifecycle, such as heterothallic fungi (with mycelial colonies of a single mating type each). Here, we investigated the evolution of recombination suppression in a clade of dung fungi encompassing 16 pseudo-homothallic and three heterothallic sibling species from the *Schizothecium* genus (Ascomycota, Sordariales). The analysis of genetic divergence based on genome sequencing indicated recombination suppression around the *MAT* locus in all investigated 13 pseudo-homothallic species. The non-recombining region ranged from 600 kb to 1.6 Mb and harbored multiple evolutionary strata, varying in size and number among species. The separation of alleles associated with alternative mating types in gene genealogies across strains within species, the high linkage disequilibrium and an inversion in one species supported the lack of recombination in the *MAT*-proximal region in pseudo-homothallic species. The overall lack of trans-specific polymorphism suggested multiple independent events of recombination suppression or the occurrence of rare events of recombination or genic conversion. Progeny analyses showed the occurrence of recombination close to the *MAT* locus in heterothallic strains. We thus revealed here multiple and likely independent evolutionary strata, associated with an extended diploid-like stage in *Schizothecium* fungi, which provides a good model for research on sex-related chromosome evolution.

## Introduction

Recombination between homologous chromosomes during meiosis is commonly thought to be advantageous in the long term as it increases the efficiency of natural selection (Rice 2002). When beneficial mutations appear at different loci in different individuals, recombination can allow them to be united on the same chromosome and thereby be both selected for (Hill and Robertson 1966). Recombination further allows preventing the accumulation of deleterious mutations in the genome by decoupling allelic associations and thereby regenerating chromosomes free of deleterious mutations from different homologous chromosomes with different deleterious mutations (Muller 1932). In some instances, however, some genomic fragments stop recombining. This phenomenon has been observed in sex-determining chromosomes in a variety of organisms, including vertebrates, insects and plants (Charlesworth 2017). Recombination suppression firstly evolves around the sex-determining locus, and can then spread to a varying extent. The spread of recombination suppression can happen in successive steps, which generates evolutionary strata, *i.e*., adjacent segments with different levels of differentiation between sex chromosomes (Charlesworth 2017). The main evolutionary hypothesis to explain the spread of recombination suppression has long been that it successively links the sex-determining gene to multiple genes under sexually antagonistic selection, *i.e.* with alleles that are beneficial in one sex and detrimental in the other one (Charlesworth 2017).

However, this hypothesis cannot explain the stepwise recombination suppression observed in fungal mating–type chromosomes, as these determine mating compatibility between individuals, but are not associated to male or female functions, or other different life-history traits (Bazzicalupo et al. 2019). Stepwise recombination suppression on mating-type chromosomes has been documented in three families of the Sordariales order in the Ascomycota, *i.e*. in the *Neurospora*, *Podospora* and *Schizothecium* genera, phylogenetically distant one from each other (Raju and Perkins 1994), and also in Basidiomycota, in the *Microbotryum, Malassezia, Ustilago, Cryptococcus, Puccinia* and *Agaricus* genera (Bakkeren and Kronstad 1994; Menkis et al. 2008; Grognet et al. 2014; Branco et al. 2018; Foulongne-Oriol et al. 2021; Coelho et al. 2023; Luo et al. 2024; Coelho et al. 2025). As of today, evolutionary strata around mating-type loci have only been reported in fungi with biallelic mating-type loci and prolonged diploid-like phases (Hartmann et al. 2021b; Jay et al. 2024), which is consistent with evolutionary hypotheses based on the sheltering of deleterious mutations ((Hartmann et al. 2021b; Jay et al. 2024). The existence of a dikaryotic phase (with two unfused nuclei of opposite mating types per cell) characterizes the Dikarya, which encompasses the two fungal orders Ascomycota and Basidiomycota. In Ascomycota, the dikaryotic phase is generally brief, karyogamy occurs almost immediately after syngamy, and mating-type is controlled by a single locus, called the *MAT* locus with two alleles (*MAT1-1* and *MAT1-2*) (Yadav et al. 2023). However, a prolonged dikaryotic or heterokaryotic life cycle, with nuclei of opposite mating types in each cell, has evolved multiple times independently in the Ascomycota, and in particular in the Sordariales order, which includes several hundreds of species distributed in different families that are frequently found in soil or dung (Marin-Felix and Miller 2022; Hensen et al. 2023). Studying additional fungal species in this order gives the unique opportunity to further test if recombination suppression evolved multiple times, in a stepwise manner and in association with a main heterokaryotic phase.

A prolonged heterokaryotic life cycle in Ascomycota is associated with pseudo-homothallism, as this breeding system corresponds to the production of self-fertile thalli, thus mimicking homothallism. While homothallic species are homokaryotic and with no incompatibility system controlling mating, pseudo-homothallic species are indeed heterokaryotic, *i.e*., with nuclei of opposite mating types (*MAT1-1* and *MAT1-2*) in hyphae. This situation contrasts with heterothallism, in which sexual reproduction can only occur by mating between different thalli, as their cells are homokaryotic and of a single mating type. In all systems, meiosis leads to sexual ascospores being arranged in ascii. In heterothallic species, ascii often contain eight ascospores, each carrying a single nucleus. In pseudo-homothallic species, in contrast, most ascii contain four ascospores, each with two nuclei of opposite mating types (Raju and Perkins 1994; Billiard et al. 2012). These two nuclei are products of a single meiosis followed by a post-meiotic mitosis and the mycelium resulting from their growth most often undergo selfing. The mating system in these fungi is thus mainly automixis (intra-tetrad mating), although they can sometimes outcross (Hartmann et al. 2021a; Ament-Velásquez et al. 2022; Vittorelli et al. 2023).

Recombination suppression and evolutionary strata can be detected in these automictic fungi based on levels of genetic divergence or heterozygosity: as automixis leads to genome-wide homozygosity, the only heterozygous regions are typically those without recombination around the *MAT* locus (Hartmann et al. 2021a; Vittorelli et al. 2023). In addition, synonymous divergence and levels of heterozygosity increase with the time since recombination suppression, as substitutions accumulate with time and remain associated with the mating type near which they appeared (Hartmann et al. 2021b); this allows detecting the successive steps of recombination suppression as “evolutionary strata” of different levels of divergence. Furthermore, recombination suppression keeps substitutions associated with the same mating type across time, which may lead to trans-specific polymorphism when recombination suppression predated speciation events, with alleles in genes from the *MAT*-proximal region clustering by mating type rather than by species (Branco et al. 2018; Hartmann et al. 2021a). In addition, crosses and analyses of progeny can be performed to assess the presence of recombination in the *MAT*-proximal region.

Regions of recombination suppression around the *MAT* locus have been identified and well studied in both *Neurospora tetrasperma* (*Sordariaceae*) and *Podospora anserina* (*Podosporaceae*), which are each complexes of closely related pseudo-homothallic species (Menkis et al. 2008; Grognet et al. 2014; Sun et al. 2017; Hartmann et al. 2021a; Hartmann et al. 2021b). In the *N. tetrasperma* species complex, a large region without recombination (*ca*. 7 Mb), spanning over more than 75% of the mating-type chromosome, links the centromere and the *MAT* locus. In *P. anserina*, the size of the region without recombination is about 800 kb, and a single obligate cross-over occurs at each meiosis between this region and the centromere. The size of the non-recombining region varies among and within species of the two species complexes (Menkis et al. 2008; Grognet et al. 2014; Hartmann et al. 2021a). Evolutionary strata have been detected in one species in the *P. anserina* complex, *Podospora pseudocomata* (Hartmann et al. 2021a), and in several lineages of the *N. tetrasperma* species complex (Menkis et al. 2008), demonstrating stepwise extension of recombination suppression in some species of these complexes. Little or no trans-specific polymorphism has been observed in the non-recombining regions, suggesting multiple independent events of recombination suppression in each of these species complexes (Menkis et al. 2008; Hartmann et al. 2021a).

Recombination suppression around the *MAT* locus has recently been reported in a third family of Sordariales, Schizotheciaceae, in the *Schizothecium tetrasporum sensu stricto* heterokaryotic CBS815.71 strain (Vittorelli et al. 2023). Non-zero synonymous divergence (d_S_) across a 1.5 Mb region around the *MAT* locus indicated suppressed recombination. Recombination suppression was confirmed by progeny analyses over 1.2 Mb. As in *P. anserina,* a single crossover occurs at each meiosis between this region and the centromere. Three evolutionary strata were distinguished based on different levels of synonymous divergence in this region, suggesting a stepwise extension of recombination suppression. The age of recombination suppression in *S. tetrasporum sensu stricto* was estimated between 40 kiloyears (ky) and 730 ky for the oldest stratum, and between 9 ky and 170 ky for the youngest stratum (Vittorelli et al. 2023; Guyot et al. 2025). The *MAT* locus in the type strain of *S. tetrasporum sensu stricto* had the typical organization of ascomycetes, with three genes in the *MAT1-1* allele and one gene in the *MAT1-2* allele, except that it harboured a *MAT1-1-1* pseudogene in the *MAT1-2* allele (Vittorelli et al. 2023). Recently, a high diversity has been revealed within the morphospecies *S. tetrasporum*, with 18 cryptic species distributed into three species complexes (*S. tetrasporum*, *S. pseudotetrasporum* and *S. octosporum,* Figure 1) (De Filippo et al. 2025). Among the three *Schizothecium* species complexes, one is heterothallic (*S. octosporum)* and two are pseudo-homothallic, these two latter being not sister groups (Figure 1). The species clusters were estimated to have diverged 10 to 15 million years ago, and species within clusters about 5 million years ago (De Filippo et al. 2025). This suggests mating-system transitions in these fungi, making them highly interesting models to test the association between pseudo-homothallism and recombination suppression.

**Figure 1:**
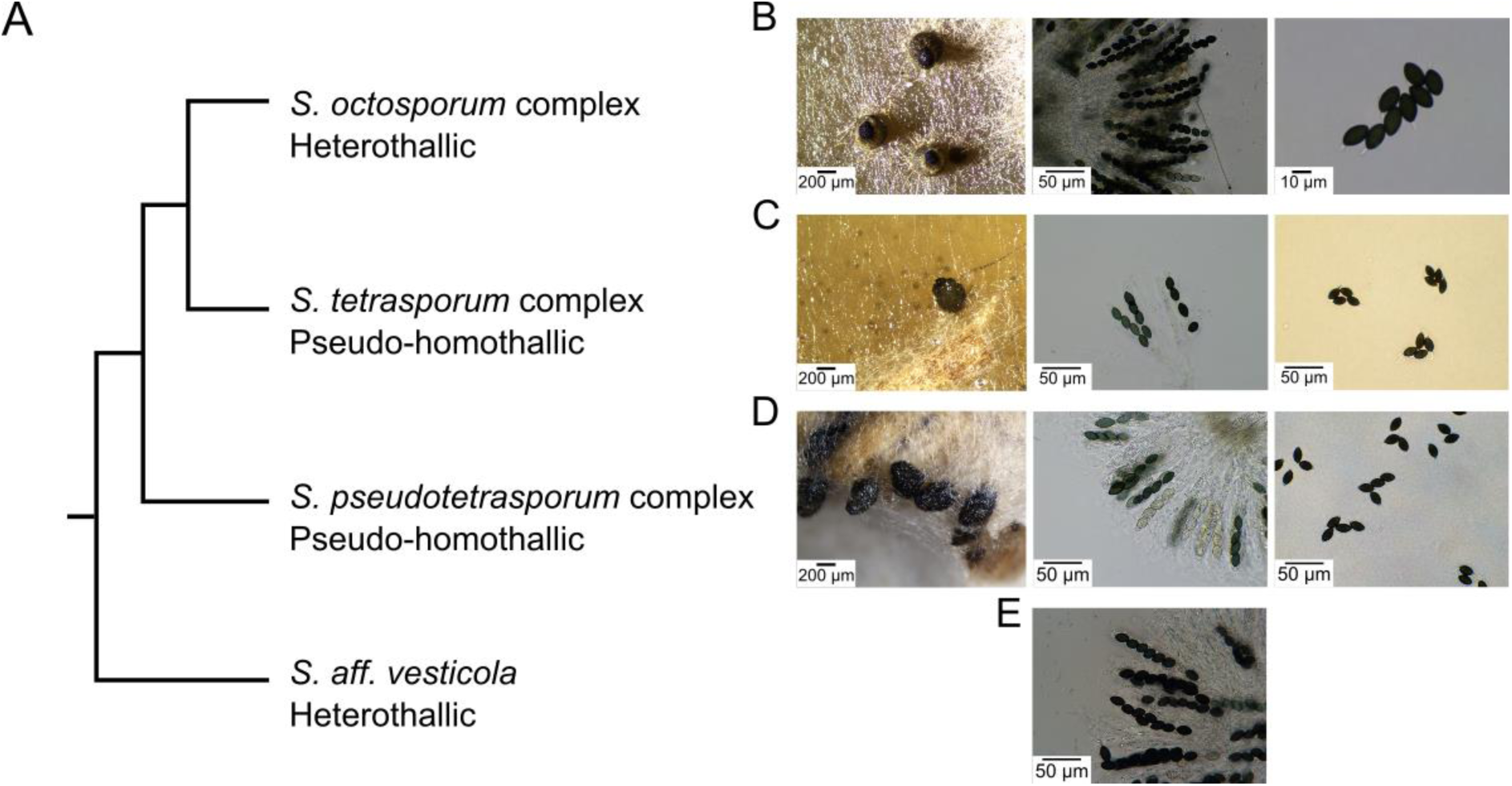
Phylogeny and pictures of the *Schizothecium species* complexes. One strain was used for each species complex. A. Cladogram of phylogenetic relationships among complexes (adapted from De Filippo et al. (2025). B. PSN1057 *Schizothecium dioctosporum*. From left to right: perithecia on M0+miscanthus media (Silar 2020), rosette of asci, spore octad. C. CBS815.71 *Schizothecium tetrasporum sensu stricto.* From left to right: perithecium on V8 medium (Silar 2020), ascii filled with spores, spore tetrad. D. PSN377 *Schizothecium pentapseudotetrasporu*m. From left to right: perithecium on V8 medium, ascii filled with spores, spore tetrad. E: PSN986 *Schizothecium aff. vesticola*. Rosette of ascii, spore octad.

While inversions between sex chromosomes are often considered as being the mechanistic cause of recombination suppression, many fungal mating-type chromosomes are entirely colinear between the two haplotypes in non-recombining regions. In the *P. anserina s*pecies complex, in several lineages of *N. tetrasperma* and in the *Schizothecium tetrasporum sensu stricto* CBS815.71 strain, the non-recombining region displays no rearrangement between mating-type chromosomes (Grognet et al. 2014; Sun et al. 2017; Vittorelli et al. 2023). Mere sequence divergence does not seem sufficient to impair recombination in *P. anserina*, and the proximal cause of recombination suppression remains unknown (Grognet et al. 2025). Inversions were identified between mating-type chromosomes in some lineages of *N. tetrasperma*, but were likely a consequence rather than a cause of recombination suppression (Sun et al. 2017). Few signatures of degeneration and no enrichment in transposable elements were detected in the mating-type chromosomes of *P. anserina* and *S. tetrasporum sensu stricto* (Hartmann et al. 2021a; Vittorelli et al. 2023).

Using available whole genomes, sequenced with short-read, from 33 heterokaryotic pseudo-homothallic *S. tetrasporum* and *S. pseudotetrasporum* strains, belonging to 13 species (De Filippo et al. 2025; Guyot et al. 2025), we scanned the mating-type chromosome for heterozygosity, to look for footprints of recombination suppression and of evolutionary strata. We also built genealogies for genes in the *MAT*-proximal region to investigate the separation of alleles associated with opposite mating types within species, as well as trans-specific polymorphism, this latter to test whether recombination suppression events occurred before or after speciation events. We further generated eight new high-quality long-read genome assemblies from the three species complexes (*S. tetrasporum*, *S. pseudotetrasporum* and *S. octosporum,* Figure 1) and their closest outgroup. We used the long-read assembly of two heterothallic strains and progeny analyses to assess the occurrence of recombination around the *MAT* locus, and thereby test for an association between pseudo-homothallism and recombination suppression. We also investigated the *MAT* locus gene organization, as well as the genomic rearrangements in the *MAT*-proximal region between mating-type chromosomes and between species.

## Results

### Recombination suppression in pseudo-homothallic lineages

Using whole genome sequencing data of 33 strains from 13 pseudo-homothallic species of the *S. tetrasporum* and *S. pseudotetrasporum* species complexes (De Filippo et al. 2025; Guyot et al. 2025) (Figure 1; Supplementary Table S1), we investigated nucleotide divergence patterns between mating-type chromosomes to detect footprints of recombination suppression and evolutionary strata in the *MAT*-proximal region. For this goal, we mapped reads on the CBS815.71-sp3 homokaryotic *MAT1-2* assembly (*i.e*., the type strain, belonging to the *S. tetrasporum sensu stricto* species) and we called SNPs. Genome coverage ranged from 16X to 168X, with an average of 72X. Because pseudo-homothallic Sordariales fungi display high selfing rates, their genomes are typically homozygous, except in regions where recombination is suppressed around the *MAT* locus. We therefore computed the density of heterozygous synonymous SNPs for each gene (*i.e*., mean number of heterozygous SNPs per synonymous site), based on the SNP calling. We will hereafter call this density of heterozygous synonymous SNPs the heterozygosity score and will use it to detect regions of recombination suppression.

The presence of a heterozygous region over at least 640 kb around the *MAT* locus, contrasting with otherwise genome-wide homozygosity (Figure 2A), strongly suggests the existence of recombination suppression in the *MAT*-proximal region in all 33 pseudo-homothallic species. The size of the heterozygous region varied between strains and species, ranging from 640 kb in the CBS262.70 *S. ditetrasporum* strain to 2.45 Mb in the PSN831 *S. tetratetrasporum* strain (Figure 2A). The size variation of the heterozygous region was largest on one side of the *MAT* locus, itself 1.82 Mb from the putative centromere in the CBS815.71-sp3 reference assembly. The heterozygous region displayed a maximum of 1.76 Mb difference in length between strains in the *MAT*-proximal region on the side towards the telomere and only 0.33 Mb on the side towards the centromere (Figure 2A). The *MAT* locus was located near the border of the heterozygous region on the site towards the centromere in all strains but the CBS262.70 *S. ditetrasporum* strain (Figure 2A). A variation in the size of the heterozygous region was also observed within species, with for instance the *S. tritetrasporum* heterozygous region extending from 1.14 Mb to 1.71 Mb across strains. The heterozygous region size was more homogeneous within *S. tetrasporum sensu stricto*, ranging from 1.38 Mb to 1.51 Mb (Figure 2A). Recent outcrossing events, which are thought to happen occasionally in pseudo-homothallic Sordariales (Hartmann et al. 2021a; Ament-Velásquez et al. 2022), can reintroduce heterozygosity in the genome, including in the flanking regions of the non-recombining region around the *MAT* locus, which may partly explain the size variation of the heterozygous region.

**Figure 2:**
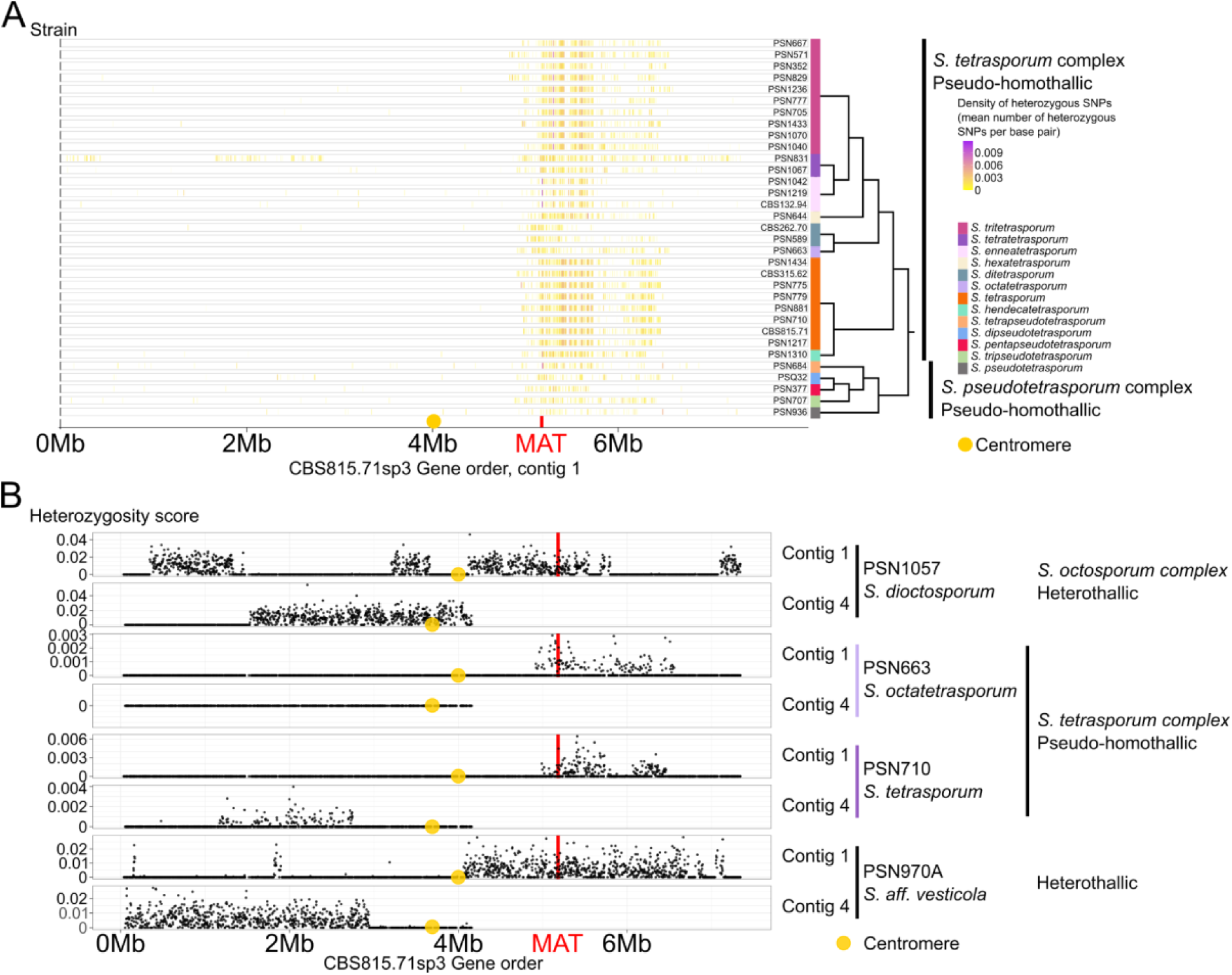
Density of heterozygous SNPs (single-nucleotide polymorphisms) in the mating-type chromosome (contig 1) and one autosome (contig 4) of strains from the *Schizothecium* species complexes. The density in heterozygous SNPs is plotted for each strain along the CBS815.71sp3 assembly. The mating-type (*MAT*) locus position is indicated by a red line and the putative centromere location with a yellow circle. A. Density of heterozygous SNPs in the mating-type chromosome of 33 strains from the *Schizothecium tetrasporum* and the *Schizothecium pseudotetrasporum* species complexes. Each vertical trait in the left part of the figure represents one gene, its color and its opacity depending on its density of heterozygous SNPs (*i.e* mean number of heterozygous SNPs per base pair for each gene). Species phylogenetic relationships inferred in De Filippo et al. (2025) are pictured by a cladogram on the right. The two species complexes are delimited by black vertical thick black lines. Species are delimited by a colored vertical rectangle. B. Density of heterozygous SNPs in the mating-type chromosome (contig 1) and one autosome (contig 4) of the CBS815.71sp3 assembly, in two pseudohomothallic strains (PSN663 and PSN710) and two heterothallic strains (PSN1057 and PSN970A). The pseudo-homothallic strain PSN710 also displays heterozygosity on the autosome contig 4. The two heterothallic strains have large regions of heterozygosity in both studied contigs. Species phylogenetic relationships inferred in De Filippo et al. (2025) are pictured by a cladogram on the right.

Inspection of heterozygosity along other contigs (hereafter referred to as autosomes) however indicated that outcrossing occurred very rarely. In order to detect footprints of recent outcrossing events, we indeed looked for heterozygous regions in autosomes in all genomes, with the criterion of at least four genes having non-null heterozygosity levels, being distant from the next gene with non-null heterozygosity by less than 50 kb. In total, only six out of 33 strains had at least one heterozygous region in autosomes (see for example strain PSN710, Figure 2B), as reported previously for four of them (Guyot et al. 2025); the other strains had no heterozygous region in autosomes (see for example strain PSN663, Figure 2B & Supplementary Figure 1A). Two of these six strains displayed heterozygous regions on more than one autosome (see for example strain PSQ32, Supplementary Figure 1C). The strain with the longest heterozygous region, PSN831 (*S. tetratetrasporum*), has undergone a very recent outcrossing event, as shown by the heterozygosity over half of its whole genome (Supplementary Figure 1B). This suggests that the variation in size of the heterozygous region across strains and species may partly be due to recent outcrossing events. However, the species with heterogeneity in the heterozygous region size, *S. tritetrasporum*, did not seem to undergo more outcrossing than the species with a homogeneous size of the heterozygous region, *S. tetrasporum sensu stricto* (one strain out of seven with heterozygosity on autosomes in *S. tetrasporum sensu stricto* and three strains out of 11 in *S. tritetrasporum*). To be conservative, we considered the strain with the shortest heterozygous region for each species as representing the non-recombining region. These strains displayed no heterozygosity on autosomes, except PSQ32 (*S. dipseudotetrasporum*) and PSN684 (*S. tetrapseudotetrasporum*) (Supplementary Figures 1C and D), that were the sole representatives of their respective species in our dataset.

The genealogies of genes located in the most heterozygous region showed that alleles clustered within species according to the mating type with which they were associated, supporting the inference of recombination suppression. Indeed, within the two species for which we had multiple genomes (*S. tetrasporum sensu stricto and S. tritetrasporum*), the alleles of 25 and 27 genes, respectively, out of the 39 genes chosen in the most heterozygous regions within the *MAT*-proximal region, including the APN2 and SLA2 genes flanking the *MAT* locus, clustered by mating type, with supported nodes, rather than by strain, for at least one mating type (Figure 3). This pattern indicates a lack of recombination, as alleles remained associated with mating types.

**Figure 3:**
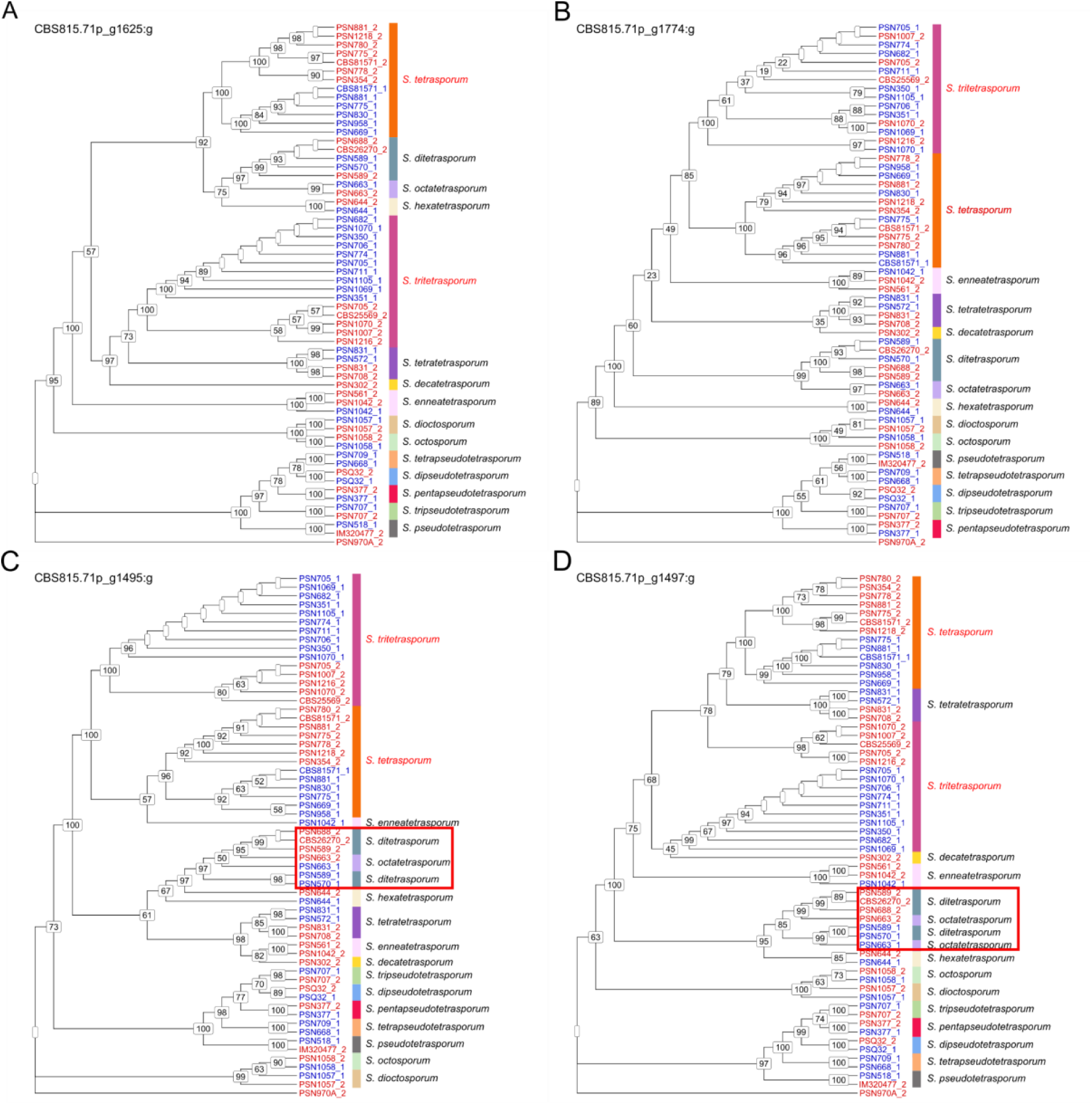
Absence of trans-specific polymorphism around the mating-type locus in the pseudo-homothallic *Schizothecium tetrasporum* and *Schizothecium pseudotetrasporum* complexes. *MAT-1-1* and *MAT1-2* mating type alleles are indicated by blue and red colors, respectively. Each Sspecies are is delimited by a colored vertical rectangle. The species *S. tetrasporum sensu stricto* and *S. tritetrasporum are writen in red*. A. The CBS815.71p_g1625:g gene with no trans-specific polymorphism but separate clustering of alleles associated with *MAT-1-1* and *MAT1-2* mating types within the species *S. tetrasporum sensu stricto* and *S. tritetrasporum*. B. The CBS815.71p_g1774:g gene with no trans-specific polymorphism and no separate clustering of alleles associated with *MAT-1-1* and *MAT1-2* mating types within the species *S. tetrasporum sensu stricto* and *S. tritetrasporum*. C. The CBS815.71p_g1495:g gene showing trans-specific polymorphism across two species (*S. ditetrasporum* and *S. octatetrasporum*), boxed in red. D. The CBS815.71p_g1497:g gene, showing trans-specific polymorphism across two species (*S. ditetrasporum* and *S. octatetrasporum*), boxed in red.

### Evolutionary strata in pseudo-homothallic lineages

We detected signatures of evolutionary strata in the heterozygous region in one strain of each of the 13 pseudo-homothallic species, *i.e*., footprints of stepwise extension of the non-recombining region (Figure 4). Evolutionary strata were identified as fragments with significantly different levels of heterozygosity scores, using a combination of methods: i) peaks in differences between mean heterozygosity scores at left and right of a sliding limit (Vittorelli et al. 2023); ii) changepoint analyses, a Bayesian inference of rupture points in series data, recently applied for the detection of evolutionary strata (Moraga et al. 2025; Rougemont et al. 2025); iii) statistical tests on the differences of heterozygosity scores between the inferred evolutionary strata (Supplementary Table S2 and Supplementary Table S3). The detection of evolutionary strata is usually performed using per-gene synonymous divergence, but this method requires separate haplotypes in the *MAT*-proximal region for the opposite mating-type chromosomes for each strain, while we only had short-read Illumina sequences for the genomes of heterokaryons for several strains, that are difficult to phase. To validate our approach based on the heterozygosity score, we compared the evolutionary strata detected with the heterozygosity score in this study in the reference CBS815.71 strain with those inferred previously using per-gene synonymous divergence (d_S_) in the same strain (Vittorelli et al. 2023). Both methods detected three evolutionary strata in the CBS815.71 strain, although boundaries were a bit shifted, by 200 kb for the youngest stratum.

**Figure 4:**
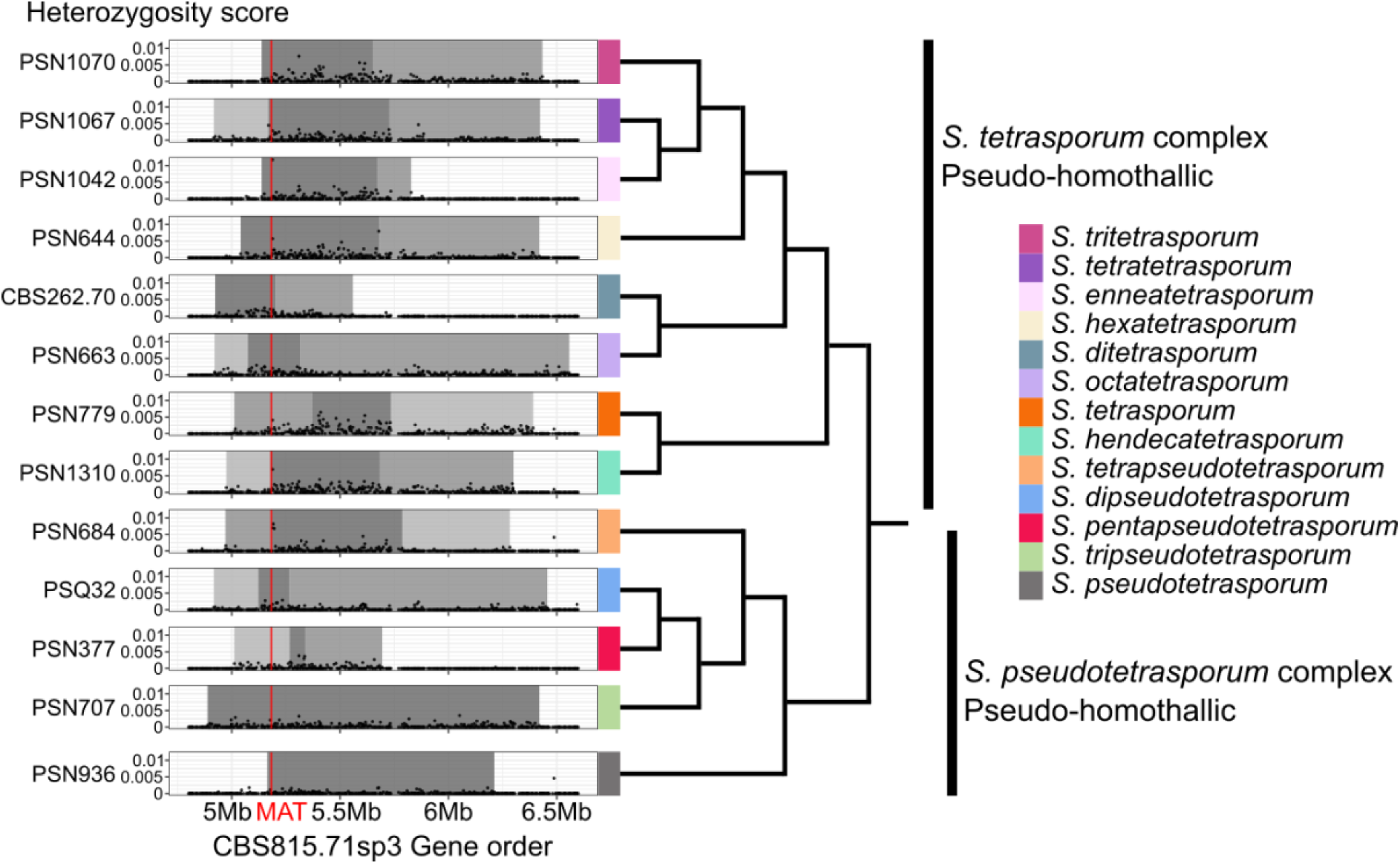
Evolutionary strata around the mating-type locus in the pseudo-homothallic *Schizothecium tetrasporum* and the *Schizothecium pseudotetrasporum* complexes. The non-recombining region around the mating-type (*MAT*) locus is represented for one strain per species along the CBS815.71sp3 assembly, represented at the bottom with the genomic scale in Megabases (Mb), and with the *MAT* locus position indicated by a red line. Each dot in the left part of the figure corresponds to one gene, and the Y axis represents the density in heterozygous SNPs (*i.e*. the mean number of heterozygous single-nucleotide polymorphisms per base pair). For each species, the strain with the shortest heterozygous region is represented on the figure. Evolutionary strata are delimited by different shades of gray, the darkest being the oldest strata, *i.e*., with the highest mean heterozygosity level. Species phylogenetic relationships are pictured by a cladogram on the right part of the figure. The two species complexes are delimited by vertical thick black lines, I being the *S. tetrasporum* cluster and II being the *S. pseudotetrasporum* cluster. The species are identified by rectangles of different colors, the same colors as in figure 1.

The number of evolutionary strata inferred in the 13 pseudo-homothallic species varied from one to three and their size and position differed among species, especially for the most recent strata (Figure 4). Two species, *S. pseudotetrasporum* and *S. tripseudotetrasporum*, both belonging to the *S. pseudotetrasporum complex,* have a single stratum, four species have two strata and seven species have three strata. The number and sizes of strata, as well as their boundaries, varied across species, within clades and even between sister species. In the seven species with three evolutionary strata, the difference in the heterozygosity score between the two youngest strata was never significant, but we nevertheless considered them as distinct strata because they were not adjacent (Figure 4): they lied at each side of the most ancient evolutionary stratum.

The *MAT* locus was located in the most heterozygous stratum in most species. In three species, however, the *MAT* locus appeared to be located outside of the most heterozygous stratum, but very close to it: at less than 4 kb in PSN684 *(S. tetrapseudotetrasporum*) and at less than 80 kb in PSN779 (*S. tetrasporum sensu stricto*) and PSN377 (*S. pentapseudotetrasporum*). The location of the *MAT* locus outside of the oldest stratum in these cases may be because the heterozygosity score is less performant to detect strata boundaries than the d_S_ index. Indeed, the mating-type locus was located outside of the most heterozygous stratum in the CBS815.71 reference strain by the change point and peak difference methods used here, although not far from it (Supplementary Figure S2), while the d_S_ approach had previously placed the *MAT* locus within the oldest stratum, also close to its boundary (Vittorelli et al. 2023).

### High linkage disequilibrium in the *MAT*-proximal region

Linkage disequilibrium further supported the lack of recombination around the *MAT* locus. We computed linkage disequilibrium on the mating-type contig for the two species for which we had the genomes for at least 10 strains (*S. tetrasporum sensu stricto* and *S. tritetrasporum*). We found a high linkage disequilibrium (LD) block for SNPs in the *MAT*-proximal region in the two species. The high-LD region spanned approximately 620 kb in *S. tetrasporum sensu stricto*, from the *MAT* locus to the extremity of the first stratum, but did not extended up to the stratum 3 (see the red triangle in Supplementary Figure 3A; mean r^2^ of all SNP pairs in the *S. tetrasporum sensu stricto* heterozygous region = 0.53, stratum 1 mean r^2^ = 0.73, stratum 2 mean r^2^ = 0.27, stratum 3 mean r^2^ = 0.07). In *S. tritetrasporum*, one of the limits of the high-LD region corresponded approximately to the *MAT* locus position. The high disequilibrium region extended over 1.08 Mb, further than the first stratum limit, covering part of the second stratum (see the red triangle in Supplementary Figure 3B; mean r^2^ of all SNP pairs in the *S. tritetrasporum* heterozygous region = 0.55, stratum 1 mean r^2^ = 0.67, stratum 2 mean r^2^ = 0.10).

### Little evidence of trans-specific polymorphism in the non-recombining regions

We found little evidence of trans-specific polymorphism in the *MAT*-proximal region, suggesting independent events of recombination suppression or frequent gene conversion events. We investigated the existence of trans-specific polymorphism in the non-recombining region by looking at gene genealogies in all pseudo-homothallic species and the two *S. octosporum* heterothallic species, using *Schizothecium aff. vesticola* PSN970A strain as outgroup, in order to assess whether recombination suppression occurred after or before speciation events in these complexes. We used only pseudo-homothallic strains sequenced as homokaryons for this analysis to have phased alleles. Considering the 39 genes *a priori* most likely to display trans-specific polymorphism (*i.e*., with heterozygosity in at least eight strains among those with the shortest heterozygous region), we found no evidence of trans-specific polymorphism in the heterozygous regions, except for two genes that showed trans-specific polymorphism for two sister species. Indeed, for 11 species and considering only well-supported nodes, the genealogies of the 39 genes analysed in the non-recombining regions branched together alleles by species and not by mating type. This suggests independent events of recombination suppression across species or occasional gene conversion events.

Only two sister species, *S. ditetrasporum* and *S. octatetrasporum,* displayed trans-specific polymorphism, at two genes, identified as CBS815.71p_g1495:g and CBS815.71p_g1497:g in the available gene model set (Vittorelli et al. 2023), located in the most heterozygous stratum, at 4 kb and 12 kb from the *MAT* locus, respectively (Figure 3). Alleles associated with opposite mating types for these two genes are indeed clustered by mating type, with supported nodes across these two species, and not by species. These two genes CBS815.71p_g1495:g and CBS815.71p_g1497:g encode proteins with functions in cell cycle (Interproscan domain IPR026000) and t-RNA linked activity (Interproscan domains IPR002300, IPR013155, IPR001569), respectively. This indicates that recombination suppression, at least at these two genes near the *MAT* locus, was ancestral to these two species.

### Recombination in heterothallic species near the mating-type locus

In contrast to pseudo-homothallic species, long genomic fragments of heterozygosity were found in heterothallic species, interspersed with long stretches of homozygosity, indicating a mixed mating system with both selfing and outcrossing (Figure 2B). Less heterozygosity was present in the PSN1057 heterothallic strain, belonging to *S. octosporum,* that branches between the two pseudo-homothallic clades, than in the PSN970A heterothallic strain, belonging to *S. aff. vesticola,* the outgroup to the three *Schizothecium* species complexes. Due to the extent of heterozygosity in these genomes as a result of outcrossing, heterozygosity is not a good proxy for recombination suppression in these heterothallic species.

We therefore investigated recombination patterns using progeny analyses from two selfing crosses of these heterothallic strains. We generated long-read-based genome assemblies for at least one mating type of these strains (PSN1057-sp6 and PSN1057-sp8 for the PSN1057 *S. dioctosporum* strain and PSN970A-sp2 for the PSN970A *S. aff. vesticola* strain) and designed three markers close to the *MAT* locus in the genomic region orthologous to the heterozygous region, in the part shared among all pseudo-homothallic species (see Material and Methods, Figure 5 and Supplementary Figure 4). In the progeny from the selfing cross from PSN1057, we found 8 recombinants out of 65 offspring between the *MAT* locus and the *H1_NRR4* marker, located at 242 kb from it, in the *MAT*-proximal region (Supplementary Figure 4). In the selfing cross of PSN970A, we found 5 recombinants out of 38 offspring between the *MAT* locus and the H2_NRR1 marker, located at 356 kb from the *MAT* locus. Among these five recombinants, one actually recombined at least as close as 86 kb from the mating-type locus, as shown by the H2_NRR3 marker located at 86 kb from the *MAT* locus. These values fit expectations for normally recombining regions in Sordariales, as three crossing-overs are expected on average per chromosome at each meiosis (Zickler et al. 1992; De Muyt et al. 2014), amounting to about 1 % recombination frequency (1 cM) every 30-50 kb. These results show that, as predicted, heterothallic strains recombine in the *MAT*-proximal region, at least as close as 242 kb from the *MAT* locus for *S. dioctosporum,* and at 86 kb from the *MAT* locus for *S. aff. vesticola*. This represents recombination events closer from the *MAT* locus than any of the recombination events found for the pseudo-homothallic strain investigated by progeny analyses in Vittorelli et al. (2023) and than the recombination suppression inferred here from genomic patterns in 13 pseudo-homothallic species (Figure 3).

**Figure 5:**
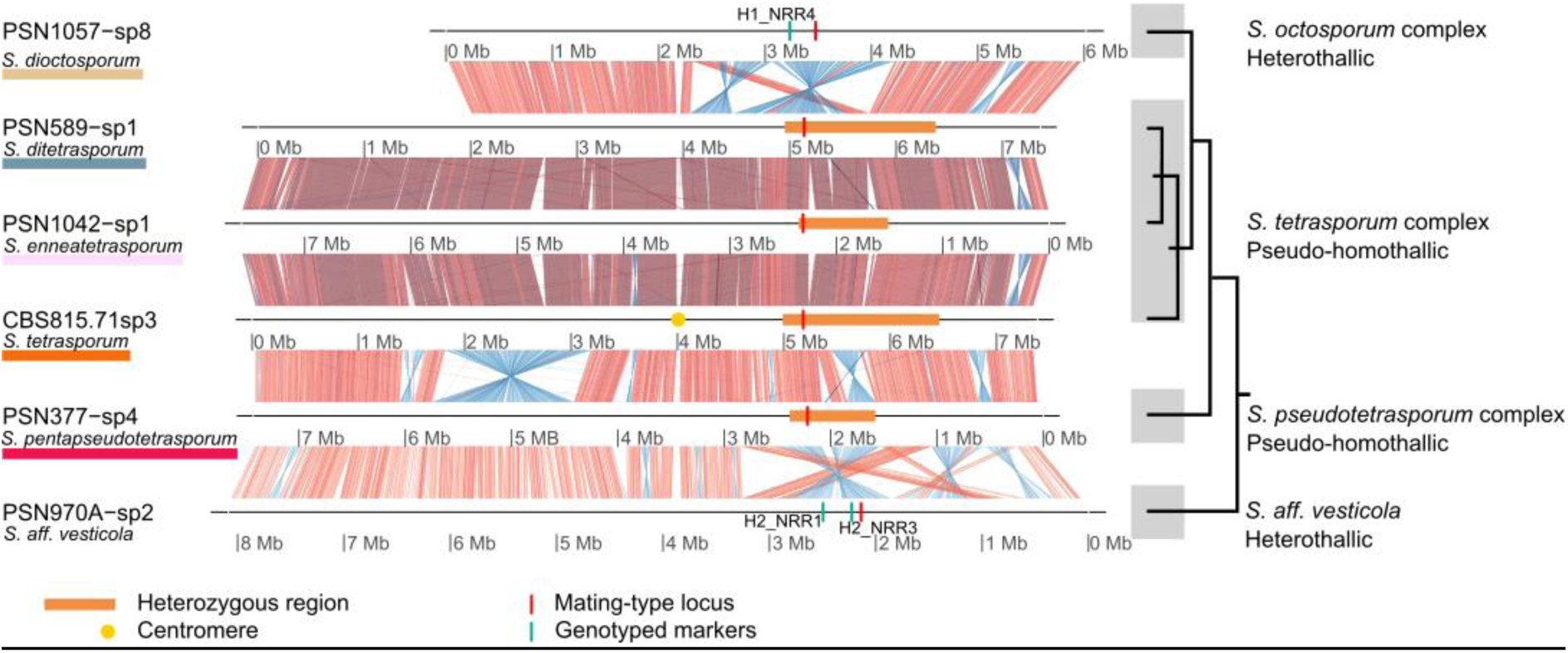
Synteny plot and genomic rearrangements between *Schizothecium species* complexes along the mating-type chromosomes. From top to bottom are shown the genome assemblies of strains PSN1057-sp8, PSN589-sp1, PSN1047-sp1, CBS815.71-sp3, PSN377-sp4 and PSN970A-sp1. Red links show collinear regions and blue links show inverted regions in the *MAT*-proximal region. The mating-type locus is located with a red bar and the putative centromere of the CBS815.71 strain with a yellow circle. The heterozygous region based on genomic divergence analyses in pseudo-homothallic strains is shown with an orange rectangle and the location of the marker tested with PCR in heterothallic strains is shown with green bars. Species phylogenetic relationships inferred in De Filippo et al. (2025) are pictured by a cladogram on the right.

### Genomic rearrangements across species and between mating types in the *MAT*-proximal region

To further explore the structure of the *MAT*-proximal region in the *Schizothecium* species complexes, we generated long-read-based genome assemblies for three additional pseudo-homothallic species (both mating types for two species), therefore obtaining eight new high-quality assemblies for the *Schizothecium* species complexes and the closest outgroup. We obtained assemblies with good BUSCO scores, with at least 95.5% of BUSCO genes present in assemblies (Supplementary Table 4).

We identified the *MAT* locus by BLAST using the *MAT1-1* and *MAT1-2* sequences of the *MAT* locus identified the CBS815.71-sp3 reference strain (Vittorelli et al. 2023) as query sequences. We found a similar organisation of the *MAT1-1* and *MAT1-2* mating-type haplotypes in all long-read assemblies of both pseudo-homothallic and heterothallic species as in the *S. tetrasporum sensu stricto* reference strain CBS815.71 (Vittorelli et al. 2023). In particular, we identified a *MAT1-1* pseudogene in all *MAT1-2* haplotypes using BLAST analyses with the *MAT1-1* pseudogene of the CBS815.71-sp3 strain as a query. The mating-type chromosome was assembled in one contig in all strains except three (PSN1057-sp6, PSN1057-sp8 and PSN377-sp3), in which it was assembled in two contigs. Telomere repeats were found at both ends of only the mating-type contig for the strain PSN377-sp4. The putative centromeric region was conserved between all sequenced strains (Figure 5).

In the two heterokaryotic strains with long-read sequencing of both mating types, the *MAT*-proximal regions of both mating types were entirely collinear in one strain and displayed an inversion in the other one (Figure 6). Opposite mating-type chromosomes were entirely collinear in the PSN377 *S. pentapseudotetraspoum* strain, as already observed in the *S. tetrasporum sensu stricto* CBS815.71 strain (Vittorelli et al. 2023). Conversely, a 430 kb inversion between the two mating-type chromosomes was present in the PSN589 *S. ditetrasporum* strain in the *MAT*-proximal region, and well-supported by long read coverage, supporting the inference of recombination suppression. The inversion covered about a third of the heterozygous region, one breakpoint roughly corresponding to an edge of the heterozygous region, with additionally 4 kb of the inversion expanding outside of the detected heterozygous region (Figure 6). As inversions prevent recombination, especially near breakpoints, we considered the non-recombining region to expand up to the end of the inversion for the following analysis.

**Figure 6:**
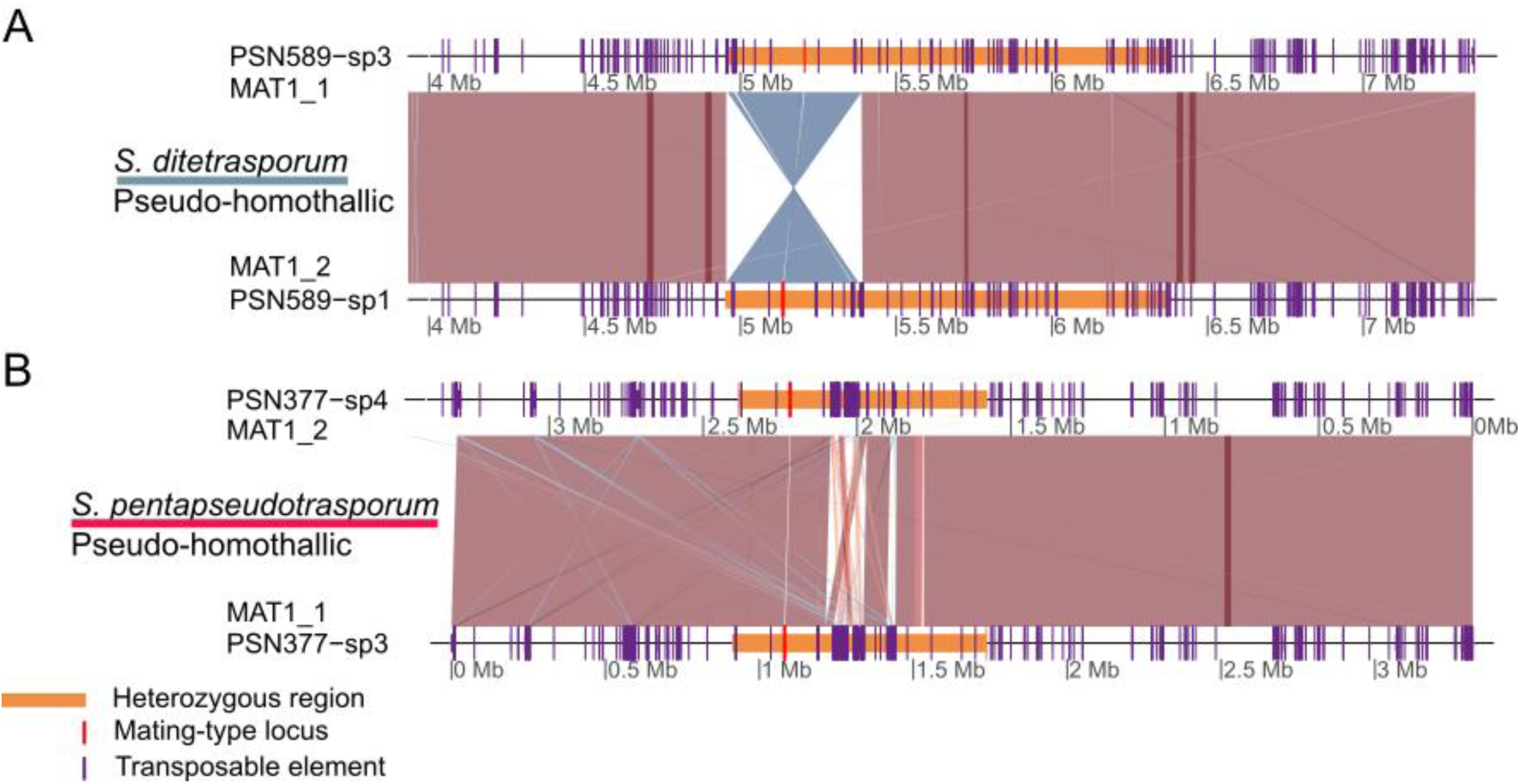
Synteny plot showing the inversion between mating-type (MAT) chromosomes of the strain PSN589 and the absence of rearrangement between mating-type chromosomes of the strain PSN377. Red links show collinear regions and blue links show inverted regions in the *MAT*-proximal region. The mating-type locus is located with a red bar. The heterozygous region is shown in an orange rectangle. Transposable elements are shown in purple.

We detected an inversion in the *MAT*-proximal region that was fixed between the two pseudo-homothallic species complexes. This 268 kb inversion was detected when comparing the high-quality genome assemblies of the PSN377-sp4 *S. pseudotetrasporum* strain one the one hand and of the three species of the *S. tetrasporum* complex (strains CBS815.71-sp3, PSN1042-sp1, PSN589-sp1) on the other hand, all having the *MAT1-2* mating type. The *MAT*-proximal region was collinear between the three strains of the *S. tetrasporum* complex. The same genomic arrangement was present in both mating types of the PSN377 *S. pseudotetrasporum* strain (PSN377-sp4 and PSN377-sp3; Figure 6). The Illumina-based genome assemblies (De Filippo et al. 2025) allowed to infer the arrangement at the inversion breakpoints for some strains, which indicated that the strains PSN707, PSQ32 and PSN684, from the S. *pseudotetrasporum* complex, had the same inversion arrangement as PSN377, from the same species complex. For one strain of the eight other species of the *S. tetrasporum* complex (PSN1070, PSN1067, PSN644, PSN663, PSN580, PSN590, PSN959, and PSN1310), we could infer collinearity at one or the two inversion breakpoints with the same inversion haplotype as the other strains of this complex sequenced with long reads, i.e., CBS815.71-sp3, PSN1042-sp1 and PSN589-sp1. The inversion therefore seems fixed between the two pseudo-homothallic species complexes. Several rearrangements (inversions and translocations) were found between the heterothallic and pseudo-homothallic species in the *MAT*-proximal region.

### No transposable element enrichment in non-recombining regions

We found no transposable element enrichment in the *MAT*-proximal non-recombining region in the three pseudo-homothallic strains for which we generated long-read assemblies (Figure 5). The percentage of base pairs occupied by transposable elements was about 7 % in the whole genome in all three strains. It was not significantly different between the non-recombining *MAT*-proximal region and the other regions of the genome, *i.e*. the autosomes and the pseudo-autosomal regions (PAR), *i.e*. the recombining regions in the mating-type chromosome. For PSN589 and PSN377 we found no significant differences among genomic compartments for the percentage of base pairs occupied by transposable elements based on Kruskal-Wallis tests on 10 kb sliding windows (PSN589-sp1: chi-squared = 2.552, p-value = 0.279, df = 2; PSN589-sp3: chi-squared = 2.412, p-value = 0.299, df = 2; PSN377-sp3: chi-squared = 0.647, p-value = 0.724, df = 2; PSN377-sp4: chi-squared = 1.382, p-value = 0.501, df = 2). In PSN1042-sp1, the percentage of base pairs occupied by transposable elements was significantly different among genomic compartments overall (Kruskall-Walis test on in 10 kb sliding windows, chi-squared = 7.760, p-value = 0.021, df = 2), but pairwise differences among genomic compartments were not significant when using Pairwise Wilcoxon tests with false discovery rate correction (*MAT*-proximal region and the autosomes, p-value = 0.194; p-value = 0.051 for the two other comparisons).

## Discussion

In this study, we found evidence of recombination suppression and evolutionary strata around the *MAT* locus in all 13 investigated pseudo-homothallic species, over a genomic region ranging from 600 kb to 1.6 Mb. The inference of recombination suppression was supported by the genomic divergence between mating-type chromosomes, the separation of alleles associated to alternative mating types in gene genealogies, high levels of linkage disequilibrium and an inversion between mating-type chromosomes in one of the species. We found variation in both the number and position of evolutionary strata between species and little evidence for trans-specific polymorphism, which suggests multiple young independent events of recombination suppression across the *Schizothecium* species complexes, or/and rare, recurrent gene conversion events. In heterothallic species, in contrast, progeny analyses showed that recombination occurred close to the *MAT* locus. Our findings thus further support the strict association found so far of recombination suppression around the *MAT* locus with pseudo-homothallism, i.e. with a heterokaryotic life cycle.

### Stepwise extension of recombination suppression in pseudo-homothallic species

We detected evidence of evolutionary strata in the pseudo-homothallic *S. tetrasporum* and *S. pseudotetrasporum* species complexes, with at least two strata in each of eleven species, out of the thirteen analysed species, indicating successive events of recombination suppression around the *MAT* locus. Notably, the number, size and position of the evolutionary strata varied among species, which supports the independence of recombination suppression extension in the various species. We also observed a variation among strains in the heterozygous-region size for some species. A part of this intra-species variation is probably due to residual heterozygosity around the *MAT* locus after recent outcrossings events, as evidenced by the presence of heterozygosity in the rest of the genome in some strains. Progeny analyses previously showed that the non-recombining region was 200 kb smaller than the heterozygous region in the strain CBS815.71 (Vittorelli et al. 2023). The observed heterozygosity may be due to outcrossing events or linkage disequilibrium at the margin of the non-recombining region (Uyenoyama 2005). However, the high-LD block was shorter than the evolutionary strata in the two species in which we had multiple genomes. This may suggest that, indeed, part of the heterozygous region is only a remnant of a recent outcrossing event. However, no LD has been observed in *P. anserina* in the non-recombining region (Hartmann et al. 2021a), despite experimental evidence of recombination suppression in progenies; this has been interpreted as a result of occasional, very rare recombination or gene conversion events. Furthermore, all strains from the *S. tetrasporum sensu stricto* species had the same limits for the heterozygous region, while this species did not seem to undergo less frequent outcrossing, as shown by the size of the heterozygous regions in autosomes. Part of the variation in size of the heterozygous region within species may therefore correspond to genuine variation in the size of the non-recombining region.

Most pseudo-homothallic *Schizothecium* species analysed here display multiple evolutionary strata, while stepwise extension of recombination suppression has been detected in a single species out of seven in the *Podospora anserina* complex (Hartmann et al. 2021a), also belonging to Sordariales. Multiple strata have also been reported in the Sordariales in multiple species of the *N. tetrasperma* species complex (Hartmann et al. 2021a; Menkis et al. 2008), as well as in the basidiomycetes, in multiple species of the *Microbotryum violaceum* species complex (Branco et al. 2018; Duhamel et al. 2022) and in *Cryptoccocus* species (Hartmann et al. 2021b; Coelho et al. 2025). In *N. tetrasperma* and *M. violaceum*, the recombination suppression is older than in *Schizothecium* or *Podospora* species, up to four millions years, and signs of degeneration are more abundant, in particular multiple inversions and transposable element load (Menkis et al. 2008; Whittle and Johannesson 2011; Whittle et al. 2011; Sun et al. 2017; Branco et al. 2018; Duhamel et al. 2023).

### Multiple independent events of recombination suppression in the *MAT*-proximal region

Several lines of evidence support the occurrence of multiple independent events of recombination suppression in the *MAT*-proximal region in the *Schizothecium* pseudo-homothallic species. The discordant limits of evolutionary strata and the lack of trans-specific polymorphism indicate the occurrence of multiple, independent events of recombination suppression across the various species of the *S. tetrasporum* species complex. The presence of heterothallic species with recombining mating-type chromosomes nested within a clade of pseudo-homothallic species with recombination suppression also supports independent events of recombination suppression in pseudo-homothallic species. The young estimated age of recombination suppression compared to the speciation dates further supports independent events of recombination suppression in the *S. tetrasporum* species complex. Recombination suppression indeed seems much younger (between 40 and 730 ky estimated for the youngest and oldest evolutionary strata in the CBS815.71 strain (Guyot et al. 2025) than the estimated speciation ages, the divergence between *S. tetrasporum* and *S. pseudotetrasporum* complexes being at least 10 My old, while the intra complex divergence has been estimated to be around 5 My old (De Filippo et al. 2025).

Furthermore, the presence of a fixed inversion between the two pseudo-homothallic species complexes, with opposite mating-type chromosomes being in contrast collinear in most species, reinforces the hypothesis of independent recombination suppression events. Indeed, if recombination was already suppressed when an inversion occurred in one mating-type chromosome after speciation, the inversion should have been restricted to this mating-type chromosome, being unable to be transferred on the alternative mating-type chromosome without recombination. Gene conversion is indeed unlikely across such a large fragment (268 kb).

Finally, the overall lack of trans-specific polymorphism supports the occurrence of independent recombination suppression events. We observed trans-specific polymorphism only at two genes out of 39, and only for two species, suggesting a likely shared recombination suppression event only between the two sister species *S. ditetrasporum* and *S. octatetrasporum,* and multiple independent recombination suppression events in other species.

Alternatively, recombination suppression may have evolved before speciation within each complex, and gene conversion or very rare recombination events could regularly reset genetic divergence between mating-type chromosomes, as suggested in *P. anserina* (Hartmann et al. 2021a; Contamine et al. 1996) and in frog sex chromosomes (Rodrigues et al. 2018).

### Association between recombination suppression and pseudo-homothallism

Our study suggests an association between pseudo-homothallism and recombination suppression in the *Schizothecium* species complexes, as already reported in other fungi (Menkis et al. 2008; Grognet et al. 2014; Hartmann et al. 2021a; Jay et al. 2024). In the *Neurospora* genus, recombination suppression evolved in pseudo-homothallic *N. tetrasperma* lineages, but not in the closest outgroup, *N. crassa,* that is heterothallic. In basidiomycete fungi, recombination suppression has only been found in species with bi-allelic *MAT* loci and an automictic mating system, and without prolonged homokaryotic stages in their life cycles (Foulongne-Oriol et al. 2021; Branco et al. 2018). The heterokaryotic life cycle may allow the sheltering of deleterious mutations and thereby trigger recombination suppression evolution (Jay et al. 2024). The presence of a sheltered load, *i.e*. recessive deleterious mutations at the heterozygous state in or near non-recombining regions, was experimentally shown to be associated to specific mating-type alleles in *P. anserina* and two *Schizothecium species, S. tetrasporum sensu stricto* and *S. tritetrasporum* (Guyot et al. 2025). A faster growth of heterokaryons compared to one of the homokaryons was indeed observed in *P. anserina*, *S. tetrasporum sensu stricto* and *S. tritetrasporum*, under different conditions. In addition, analyses of non-synonymous substitutions further supported the existence of deleterious mutations in the non-recombining region, with heterozygous missense mutations, gains of stop codons and losses of start codons (Grognet et al. 2014; Guyot et al. 2025). Gene losses have also been detected just at the margin of the non-recombining region of the reference strain of *S. tetrasporum* (Vittorelli et al. 2023).

Under the hypothesis of deleterious mutation sheltering, recombination suppression would be an evolutionary consequence of the extended diploid-like stage, itself a consequence of pseudo-homothallism or automixis (Jay et al. 2024). Pseudo-homothallism may thus have evolved earlier than recombination suppression, and even perhaps at the basis of the *Schizothecium* clade studied here, with a reversion to heterothallism in the *S. octosporum* lineage. Alternatively, pseudo-homothallism may also have evolved multiple times independently in the *Schizothecium* fungi, although this hypothesis seems less parsimonious than a single origin at the basis of the clade, especially as the feature of dikaryotic spores with opposite mating types may be complex to evolve.

In *S. tetrasporum sensu stricto*, as in *P. anserina*, a single systematic crossing over occurs between the centromere and the *MAT* locus, as indicated by the high frequency of second-division segregation of the *MAT* locus (Vittorelli et al. 2023). This single crossing-over places haploid nuclei of opposite mating types next to each other after meiosis in the tetrad to form self-fertile heterokaryotic ascospores, being thus essential for pseudo-homothallism. It remains to be studied if this single crossing over occurs also in other pseudo-homothallic species of the *S. tetrasporum and S. pseudotrasporum* complexes. As discussed in (Vittorelli et al. 2023; Grognet et al. 2025). An unknown mechanism compacting two nuclei per ascospore, for the production of self-fertile ascospores, also evolved. Whether recombination suppression is a cause or a consequence, or acts in synergy with these mechanisms necessary for pseudo-homothallism, remains to be studied. Despite the heterogeneity in size and position of evolutionary strata across pseudo-homothallic species, a shared feature was that the non-recombining region always expanded in the same direction, systematically away from the centromere, toward the chromosome arm end. The lack of extension of recombination suppression closer to the centromere may be caused by constraints due to the single and systematic crossing over occuring between the *MAT* locus and the centromere in this species.

Under this hypothesis, recombination suppression should extend starting from the MAT locus. It was therefore surprising to find the *MAT* locus outside of the most ancient evolutionary stratum in some strains. However, this may just be due to imprecise delimitation of stratum boundaries. Delimiting strata is indeed notoriously difficult, especially when gene conversion can occur and reset genetic divergence in some genes.

### Conclusion: a new model for studying young sex-related chromosomes

In conclusion, we reveal here multiple events of recombination suppression along mating-type chromosomes and across closely related species, and restricted to species with a main diploid-like phase. This adds strong support to the view that recombination suppression often evolves and extends stepwise around the *MAT* locus in fungi despite the lack of sexual antagonism, but only in species with a diploid-like life cycle. This informs on the mechanisms selecting for recombination suppression, being consistent with the sheltering of deleterious mutations (Jay et al. 2024). Furthermore, the recombination suppression events are young in the *Schizothecium* complex, providing a new model for studying the first steps of recombination suppression and its consequences.

## Material and Methods

### Strain origins

We studied a total of 67 fungal strains, including 55 pseudo-homothallic strains from the *S. tetrasporum* complex, nine pseudo-homothallic from the *S. pseudotetrasporum* complex, two heterothallic strains from the *S. octosporum* complex and one strain from the heterothallic sisters species *Schizothecium aff. vesticola*. A robust phylogeny and taxonomic description of these strains have been reported in De Filippo et al. (2025). Fifty-eight strains are kept frozen at –80°C at the Laboratoire Interdisciplinaire des Energies de Demain, University Paris-Cité and at the Ecologie, Société et Evolution, University Paris Saclay, France labs and are available upon request. Five strains are available at the Westerdijk Fungal Biodiversity Institute, the Netherlands and one at the CABI culture collection. Table S1 gives information on strain origin and species.

### Genomic data

We used whole genomes of fungal strains available in Genbank under the BioProject accession number PRJNA1190690 that were sequenced with the Illumina technology, either as heterokaryons or homokaryons (De Filippo et al. 2025) (see Supplementary Table S1 for accession numbers of raw reads and genome assembly). Briefly, homokaryotic spores can be obtained during strain isolation when five-spores asci filled with three heterokaryotic spores and two homokaryotic spores of opposite mating type are present. When no five-spores asci were found during isolation, we produced a F1 generation by selfing. We isolated homokaryotic spores of opposite mating types from five-spore asci expelled from perithecia and determined by PCR screening the mating type allele using previously designed primers (Vittorelli et al. 2023). We extracted DNA with the Nucleospin soil kit from Macherey Nagel. Library preparation and Illumina genome sequencing were performed in six different batches by Genewiz, Genoscope or Novogene as described in (De Filippo et al. 2025). We further used as reference the long-read-based genome assemblies of the strains CBS815.71-sp3 and CBS815.71-sp6 available from the Genbank BioProject accession number PRJNA882797 at assembly accession number JAQKAE000000000 for CBS815.71-sp3 and JAQKAD000000000 for CBS815.71-sp6 (Vittorelli et al. 2023).

### Mapping and SNP calling

We first checked Illumina raw read quality using FastQC v0.11.1 v.0.11.5 (Andrews 2010), except for the Genoscope batch as FastQC reports were supplied by the platform. Reads were trimmed with Trimmomatic v0.36 (Bolger et al. 2014) in paired-end mode with the TruSeq3 adapter and the following options: PE –threads 2 –phred33 TruSeq3-PE.fa:2:30:10 LEADING:10 TRAILING:10 SLIDINGWINDOW:5:10 MINLEN:50. The genomes sequenced by the Genoscope were already trimmed. Mapping and SNP calling for the 64 studied strains were performed using the pipeline described in De Filippo et al. (2025). Illumina trimmed reads were mapped against the CBS815.71-sp3 (*MAT1-2*) long-read assembly (Vittorelli et al. 2023) with bowtie2 v2.3.4.1 (Langmead et al. 2009) with the options –very-sensitive-local –phred33 – X 1000. We used the MarkDuplicates tool from PicardCommandLine v2.8.1 to get rid of duplicated sequences. The mating types of the sequenced genomes were checked by assessing visually if the coverage at the *MAT* locus was >10X with Integrative Genomics Viewer v2.15.1 in the mapping on the reference genomes of the two mating types, CBS815.71-sp3 and CBS815.71-sp6 (Vittorelli et al. 2023). To do so, heterokaryons were also mapped against the CBS815-71sp6 (*MAT1-1*) long-read assembly (Vittorelli et al. 2023), to check the presence of the *MAT1-1* allele, as possibly unbalanced ratios of *MAT1-1* and *MAT1-2* nuclei or nucleus loss may lead to unbalanced coverage of the *MAT* locus region. We called SNPs against the CBS815.71-sp3 assembly with the HaplotypeCaller tool of the Genome Analysis Toolkit (GATK) v4.1.2.0 (McKenna et al. 2010) in the diploid mode. We used CombineGVCF, GenotypeGVCF and SelectVariants to merge data from all strains and conjointly call SNPs. GATK Good Practice rules were followed with some adaptations to filter variants for quality with the following options: QD=20.0; FS=60.0; MQ=20.0; MQRankSumNeg=-2.0; MQRankSumPos=2.0; <SOR=3.0; QUAL=100. SelectsVariants and VCFtools v0.1.17 were used to filter out indels, and keep only good-quality SNPs with the 90% missingness option. We kept only biallelic SNPs, polymorphic within our pool of strains.

### Detection of evolutionary strata

We investigated recombination suppression around the *MAT* locus in *S. tetrasporum* strains based on sequence divergence between mating-type chromosomes. As pseudo-homothallic fungi generally reproduce mostly by selfing, their genomes are indeed highly homozygous, except in the genomic regions without recombination capturing the permanently heterozygous *MAT* locus (Hartmann et al. 2021a, Vittorelli et al. 2023). We computed a heterozygosity score to detect recombination suppression using heterozygous SNPs along the genome in heterokaryons and SNPs occuring between homokaryons of opposite mating types but originating from the same heterokaryon. Strains sequenced as heterokaryons were used in this analysis if the two mating-type regions had a coverage of at least 10X each, as checked with the genomecov tool of bedtools (v2.26.0) (Quinlan and Hall 2010). If a mating-type haplotype had a coverage of less than 1X, the strain was considered as a homokaryon and included in the analysis only when the opposite mating type was available. Homokaryotic strains were used in this analysis when two homokaryons of different mating types were sequenced for the same isolated heterokaryotic strain. We selected the SNPs located in genes using bedtools (v2.26.0) (Quinlan and Hall 2010) and the previously published *S. tetrasporum sensu stricto* reference strain gene model (Vittorelli et al. 2023). We obtained 2,525,078 genic SNPs for the 50 genomes used to compute heterozygosity (see Supplementary Table S1). We then selected synonymous SNPs with SNPeff (v5.2c) (Cingolani et al. 2012) and calculated for each gene a heterozygosity score between 0 and 1 by dividing the number of heterozygous SNP per gene by gene length (bp). Gene overlaps in the gene model led some SNPs to be counted twice. To avoid over-representation of these SNPs, we used the R metapackage tidyverse (v2.0.0) to filter out the second occurrence. We plotted the heterozygosity score along the CBS815.71sp3 assembly with the ggplot2 R package (v3.5.1) for each strain. We delimited the non-recombining region on each side by searching, in the chromosome arm carrying the *MAT* locus, the suites of at least four consecutives genes located farthest from the MAT locus with each at least one heterozygous SNP and less than 50 kb between the midpoints of two consecutive genes. We measured the size of the non-recombining region by calculating the distance between the midpoints of the farthest of these genes at the two sides of the *MAT*-proximal region. We identified additional heterozygous regions, located on the autosomes or on the mating-type chromosome arm which does not carry the MAT locus, using a customized script. We chose to define them as chromosomal regions consisting of at least four genes with at least one synonymous heterozygous SNP and less than 50kb between the midpoints of two consecutive genes.

For each species, we selected the strain with the shorter non-recombining region in order to detect evolutionary strata, which was done by combining two different methods. First, we identified putative strata limits using the method previously described (Vittorelli et al. 2023) based on the detection of peaks in the difference in the heterozygosity score on each side of a sliding limit. Second, we used the change-point method, with the mcp function from the mcp R package (v0.3.4) (Lindeløv 2020) to determine the number of strata and test the previously obtained limits. We detected locations where the distribution of the heterozygosity score changes, by setting 1, 2 and 3 change points, corresponding to evolutionary strata limits, without setting *a priori* for the change-point locations. We retained the stratum limits that co-localized between the first method and a predicted change-point in at least one of the three change-point analysis (Supplementary Figure S2A and S2B). Finally, we used a pairwise Wilcoxon test with a Benjamini & Hochberg correction (Benjamini and Hochberg 1995) from the stats (v4.4.3) R package to test for significant differences between strata in the mean synonymous heterozygosity score per bp and per gene. When two adjacent strata had a non-significant difference in mean heterozygosity, we merged them as one stratum and ran the Wilcoxon test again. We compared the results obtained by this method, based on either d _S_ (synonymous divergence) or numbers of synonymous heterozygous SNPs for the CBS815.71 strain (*S. tetrasporum sensu stricto*) (Supplementary Figure S2C).

### Analyses of trans-specific polymorphism

We built allele genealogies for genes located in the non-recombining region and searched for trans-specific polymorphism, in order to determine with recombination suppression occurred before or after speciation between the *S. tetrasporum* and *S. pseudotetrasporum* species complexes. Indeed, as soon as recombination is suppressed, mutations accumulate independently in the regions flanking the *MAT* locus and remain associated with the mating type near which they appeared. We selected the 39 genes *a priori* most likely to display trans-specific polymorphism, corresponding to all genes with at least one heterozygous synonymous SNP in at least eight strains, among those with the shortest heterozygous region in each species. We selected the 60 homokaryotic genomes, corresponding to 45 strains from eight species in the *S. tetrasporum* complex, ten from five species in the *S. pseudotetrasporum* complex, four from two species in the *S. octosporum* complex and the PSN970A *Schizothecium sp.* strain, used as an outgroup (See Supplementary Table S1), based on the whole genome phylogeny provided in (De Filippo et al. 2025). To select genic SNPs, we used the previously established *S. tetrasporum sensu stricto* reference strain gene model (Vittorelli et al. 2023) and bedtools (v2.26.0). We converted the acquired VCF file to phylip format with PGDSpider (v2.1.1.5) (Lischer and Excoffier 2012) and used IQ-TREE (v1.6.1) (Nguyen et al. 2015; Kalyaanamoorthy et al. 2017; Hoang et al. 2018) with ModelFinder to obtain a phylogeny with the best fit model. We designated PSN970Asp2 as the outgroup with the –o option and we used the –s –bb 1000 –alrt 1000 options to compute ultrafast bootstraps of a 1000 iterations. The PSN518m strain (*S. pseudotetrasporum*) was removed with VCFtools (v0.1.17) from the SNP set located in the CBS815.71p_g1623:g gene, as it contained only gaps or missing values. All trees were visualized with the ggtree package (v3.10) in R (Yu 2020; Yu et al. 2018; Yu et al. 2016). For the tree analyses, we considered that a bootstrap inferior to 90 indicated a non-resolved branch.

### Long-read sequencing, assembly and analyses

We sequenced eight genomes with Oxford Nanopore Technologies (ONT) to obtain long reads so as to get high-quality genome assemblies of the *MAT*-proximal non-recombining region. We selected three pseudo-homothallic strains, one from the *S. pseudotetrasporum* complex (PSN377, *S. pentapseudotetrasporum*) and two from the *S. tetrasporum* complex, chosen for their different non-recombining region size (PSN589, *S. ditetrasporum* and PSN1042, *S. enneatetrasporum*). We sequenced two homokaryons of opposite mating types for PSN377 (PSN377-sp3 and PSN377-sp4) and PSN589 (PSN589-sp1 and PSN589-sp2) and the *MAT1-2* homokaryon for PSN1042 (PSN1042-sp1). We also sequenced with ONT the genomes of two homokaryons of opposite mating types from the heterothallic PSN1057 strain (PSN1057-sp6 and PSN1057-sp8) and the *MAT1-2* genome of the PSN970A strain (PSN970A-sp2). Illumina whole genomes were already available for these strains and were downloaded from the GenBank database under bioproject PRJNA1190690 (see Table S1 for accession number; (De Filippo et al. 2025)). For DNA extraction, mycelium was grown in M2 liquid medium (Silar 2020) in darkness at 22°C for 10 days. Mycelium was harvested using a plastic funnel and filtration paper. Mycelium was stored at –80°C for 24 hours, and then lyophilized for 24 hours. DNA extraction of high molecular weight was performed as described previously (Vittorelli et al. 2023). Sequencing was performed *in house* using a MinION MK1c device and with the ligation sequencing kits as described in Table S4.

We assembled the raw reads with Flye (v.2.9.3-b1797n, Kolmogorov et al. 2019) setting an expected genome size of 35 Mb (Vittorelli et al. 2023). The assembly was polished using raw long reads and Illumina reads as recommended. We aligned raw ONT reads on the Flye assembly with minimap (v2.17-r941) and sorted them with samtools (v1.9). We then polished the assembly with marginPolish (v1.3.dev-5492204) with the allParams.np.microbial.r94-g305.json model. We corrected the assembly with mapping ILLUMINA reads with bwa (v0.7.17-r1188) using the command bwa mem and the following options: –t 14 –x ont2d, and we then sorted them with samtools (v1.9). We checked our assembly quality by running BUSCO (v5.5.0) before and after the correction step, using the Sordariomycota ortholog set (sordariomycetes_odb10) and run Assembly-stats (v1.0.1) and emboss (v6.6.0.0) to obtain the assembly statistics. To identify telomeres, we search for telomeric repeats “TTAGGG” and “CCCTAA”. We identified the *MAT* locus location by blast using the *MAT1-1* and *MAT1-2* sequences of the *MAT* locus identified the CBS815.71sp3 reference strain (Vittorelli et al. 2023) as query sequences.

We used nucmer (v3.1) (Marçais et al. 2018) to investigate the synteny between the long read based genomes. Output of nucmer was filtered to keep only genomes fragments with at least 70% of identity between genomes using a customized script and plotted with the R package genoPlotR (V.0.8.11) (Guy et al. 2010). To investigate synteny in the 268 kb inversion found in the *MAT*-proximal non-recombining region between the long read-based genomes of PSN377 *S. pseudotetrasporum* strain and the three species of the *S. tetrasporum* complex (see Main results), we used Illumina-based genome assemblies (De Filippo et al. 2025) for one strain of each of the twelve remaining pseudo-homothallic species. We studied the genome assemblies of four additional species of the S. *pseudotetrasporum* complex (strains PSQ32, PSN707, PSN684 and PSN936) and eight species of the S. *tetrasporum* complex (strains PSN1070, PSN1067, PSN644, PSN663, PSN580, PSN590, PSN959 and PSN1310). Genomes were downloaded from the GenBank database under bioproject PRJNA1190690 (see Table S1 for accession number). We used the tool D-GENIES (Cabanettes and Klopp 2018) to visualize synteny between the Illumina-based genome assemblies and the long-read assemblies.

We annotated transposable element content and investigated enrichment of transposable elements in the non-recombining region in a similar way as described previously (Vittorelli et al. 2023). Briefly, we run RepeatModeler v1.0.11 (Hubley et al.) on each long-read genome assembly in order to *de novo* identify consensus sequences of transposable elements. We used each set of consensus sequences with RepeatMasker v4.0.9 (Smit et al.) to annotate repeats for each assembly. We parsed the RepeatMasker outputs and removed low-complexity and simple repeats with the parseRM_-merge_interrupted.pl script from https://github.com/4ureliek/Parsing-RepeatMasker-Outputs (last accessed June 29, 2024).

We computed transposable element density in genomic windows using a customized R script. Overlapping transposable elements annotated as belonging to the same family were merged into single elements to compute transposable element density.

### Crosses and segregation analyses in heterothallic strains

We analyzed progenies in order to test for recombination suppression in two heterothallic strains, PSN1057 (*S. dioctosporum*) and PSN970A (*S. aff. vesticola*). Indeed, as the genomes of heterothallic strains are not homozygous even in recombining regions, heterozygosity cannot be used as a proxy for recombination suppression. We induced sexual reproduction in these heterothallic strains as previously described (De Filippo et al. 2025). For each strain, we crossed two F1 homokaryotic spores of opposite mating types to generate a F1 progeny and isolated about a hundred of F2 ascospores from the selfed cross of PSN1057 and from the selfed cross of PSN970A, all from different ascii. We used the same protocol as for the *S. tetrasporum sensu stricto* CBS815.71 strain to induce germination and grow the ascospores as described in (Vittorelli et al. 2023). To extract DNA of F2 ascospores a protocol based on Chelex as in (Guyot et al. 2025). We used PCR to test if recombination occurred in the heterozygous region flanking the *MAT* locus. We designed for this goal PCR primers to determine the mating type of each haploid offspring and its genotype in a region corresponding to the oldest stratum in CBS815.71. For each cross, we genotyped the mating-type allele of all F2 ascospores using two primers pairs (one pair for *MAT1-1*, one pair for *MAT1-2*) (Supplementary Table S5). We further genotype one marker at 242 kb from the mating-type locus for PSN1057 (H1_NRR4), and two markers for PSN970A, at 86 kb (H2_NRR1) and 356 kb (H2_NRR3) distance from the *MAT* locus (Supplementary Table S5). Only the offspring found to be recombining between the *MAT* locus and the H2_NRR1 marker were genotyped with the H2_NRR3 marker. We designed primers using the Primer3Plus tool (Untergasser et al. 2012) based on the genomic sequences obtained in long-read assemblies. We used two sets of PCR conditions as summarized in Suplpementary Table S5 and Supplementary Table S6.

## Acknowledgements

We thank Ricardo C. Rodríguez de la Vega for advice on genomic analyses and Fabienne Malagnac for comments on a previous version of the manuscript.

## Data Availability Statement

All raw ONT reads and assemblies newly produced in this study were deposited on the GenBank database under bioproject PRJNA1190690 (see Table S1 for accession numbers).

## Author contribution

FEH, PS and TG conceptualized the study and supervised the study. FEH, PS, TG and PG acquired funding. FEH, PS and TG collected samples. PS and VG isolated strains. EDF, EC, AS, PM, JLD and AL performed experiments of culture in vitro and molecular biology. EDF, CL, PG and FEH analyzed genomes. EDF produced figures. EDF, FEH and TG wrote the original draft. All authors edited the manuscript.

