## Supplementary material for "Stepwise recombination suppression around the mating-type locus associated with a diploid-like life cycle in *Schizothecium* fungi"

Supplementary FIGURES legend

**Supplementary Figure 1: Density of heterozygous SNPs (single-nucleotide polymorphisms) in the mating-type chromosome (contig 1) and autosomes (contig 2, 3, 4, 5, 6 and 7) of four strains from the *Schizothecium* species complexes.** A. PSN663 (*S. octatetrasporum*): no heterozygous SNPs on autosomes. B. PSN831 (*S. tetratetrasporum*). Half of the autosomes have heterozygous SNPs. C. PSQ32 (*S. dipseudotetrasporum*). Two autosomes have heterozygous SNPs, highlighted in yellow. D. PSN684 (*S. tetrapseudotetrasporum*). One autosome has heterozygous SNPs, highlighted in yellow.

**Supplementary Figure 2: Illustration of the method to detect evolutionary strata in the CBS815.71 *Schizothecium tetrasporum sensu stricto* strain.** The mating-type locus location is indicated with a red vertical line. A. Detection of evolutionary strata using the method of Vittorelli et al. (2023). The position of boundaries of detected evolutionary strata are indicated with vertical dotted lines. (1) Heterozygosity score along the CBS815.71sp3 assembly in the non-recombining region; (2), (3) and (4): Variation of the heterozygosity score when dividing the non-recombining region into two segments, with the limit sliding along genes, and the difference of heterozygosity score between the two segments computed for each partition; in sub-panel (2) the variation was computed between 5,200,000 bp and the end of the heterozygous region to delimit the two first strata; in sub-panel (3) the variation was computed between between the start of the non-recombining region and the previously detected boundary to delimit a third stratum; in sub-panel (3), the variation was computed between between the start of the non-recombining region and the second boundary detected to delimit a fourth stratum. For 5 species, no boundary could be found between the start of the non-recombining region and the second detected boundary, as they were too close together. The variation was then computed between the two first detected boundaries, to delimit a fourth stratum. B. Detection of evolutionary strata using the change-point method with one (1) two (2) or three (3) change points. The position of boundaries of detected evolutionary strata using the method of Vittorelli et al. (2023) (panel A) are indicated with vertical orange lines. C. Comparison of evolutionary strata detected using (1) the heterozygosity score by retaining the stratum limits that co-localized between the first method and a predicted change-point in at least one of the three change-point analysis and (2) $d_S$ values computed in (Vittorelli et al. 2023) by retaining the stratum limits that co-localized between the first method and a predicted change-point in at least one of the three change-point analysis.

**Supplementary Figure 3: Linkage disequilibrium (LD) along the contig carrying the mating-type locus in *Schizothecium tetrasporum sensu stricto* (A) and *Schizothecium tritetrasporum* (B).** LD heatmap using single nucleotide polymorphisms (SNPs) located on the contig carrying the mating-type locus (contig 1 of the CBS815.71-sp3 reference genome). The number of SNPs was 7,277 for the 18 studied strains in *S. tetrasporum sensu stricto* and 6,421 for the 27 studied strains (29 genomes) in *S. tritetrasporum.* The block of SNPs with high LD, i.e. $r_2$>0.9, corresponds to the red triangle. The mating-type locus location is indicated with a red bar and the region lacking recombination with an orange rectangle. The heatmap is plotted along SNP order. The total length of the contig is also indicated.

**Supplementary Figure 4: Proportion of offspring with recombination events between the mating-type locus and neighboring markers in selfed crosses of three strains,**

**belonging to two heterothallic *Schizothecium* species and a pseudo-homothallic *Schizothecium* species.** A. Recombination frequency in centimorgans (cM) around the mating-type locus in the pseudo-homothallic heterokaryotic *Schizothecium tetrasporum sensu stricto* reference CBS815.71 strain (data from Vittorelli et al. 2023). Recombination frequency was estimated here in two heterothallic species, by genotyping 107 offspring after one round of experimental selfing at the mating-type locus and at surrounding heterozygous loci (P_NRR2 and P_NRR5), whose relative positions are illustrated by red and gray vertical lines, respectively. Genomic distances (in kilobases) between the loci are displayed below the black arrows. Red, green and blue rectangles indicate the different evolutionary strata based on $d_S$ values. In the B and C panels, the number of recombinant offsprings compared to the number of tested offsprings in this study is indicated below the gray arrows. The relative locations of the mating-type locus (*MAT*) and markers tested are shown with vertical bars. Genomic distances (in kilobases) between markers are indicated below the black arrows. B. Cross PSN1057-sp6 x PSN1057-sp8. C. Cross PSN970A-sp2 x PSN970A-sp1. Only the recombinant offspring detected when genotyping the H2_NRR1 marker were then genotyped with the H2_NRR3 marker.

Supplementary Table legend

**-Table S1: Information on the fungal strains used in this study and accession numbers of their whole genome sequences.** Information was retrieved from De Filippo et al (2025).

**-Table S2: Localization of the evolutionary strata detected in the pseudo-homothallic *Schizothecium tetrasporum* and the *Schizothecium pseudotetrasporum* complexes.** Genomic coordinates are given along contig 1 of the CBS815.71-sp3 genome.

**-Table S3: Results of statistical tests for assessing the significance of differences in number of heterozygous SNPs between evolutionary strata in the pseudo-homothallic *Schizothecium tetrasporum* and the *Schizothecium pseudotetrasporum* complexes.**

**-Table S4: Summary statistics of raw reads and genome assemblies for the five *Schizothecium* strains sequenced with the long-read technology Oxford Nanopore.**

**-Table S5: Primers used to test recombination suppression in the MAT-proximal region.**

**-Table S6: PCR conditions used to test for recombination suppression in the MAT-proximal region.**

Supplementary Figure 1:

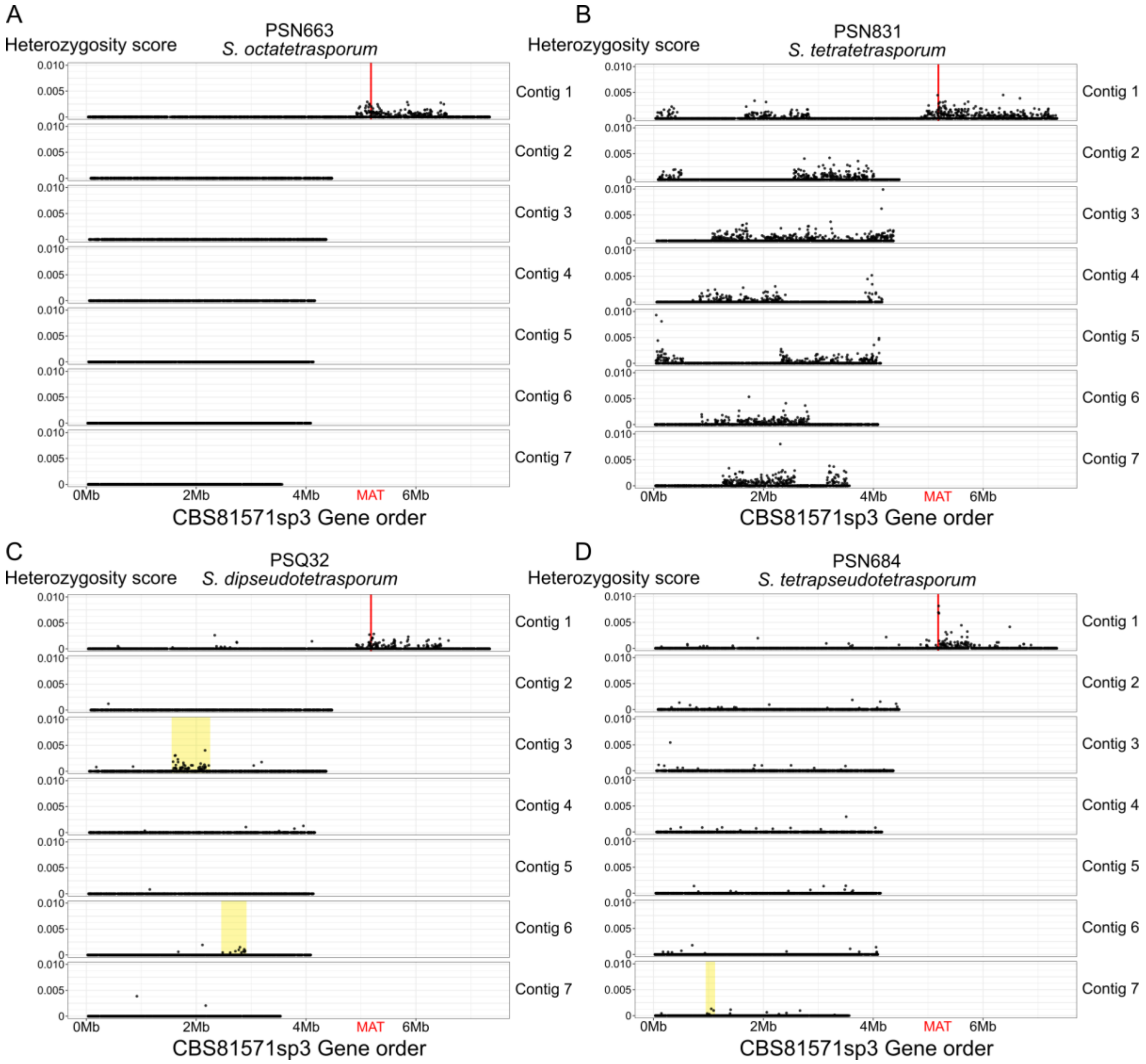

Supplementary Figure 2:

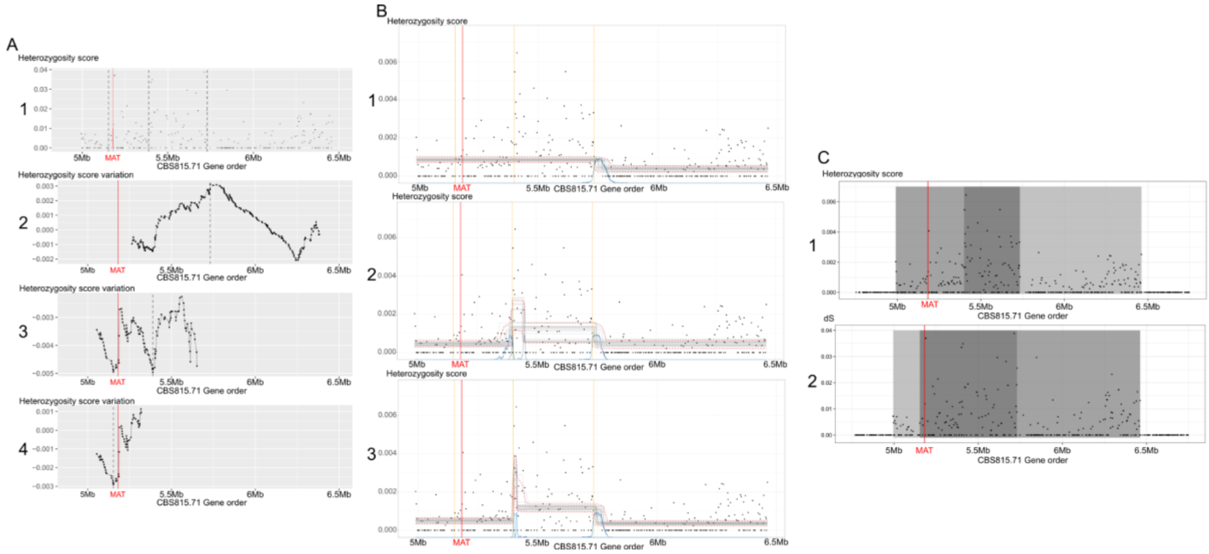

Supplementary Figure 3:

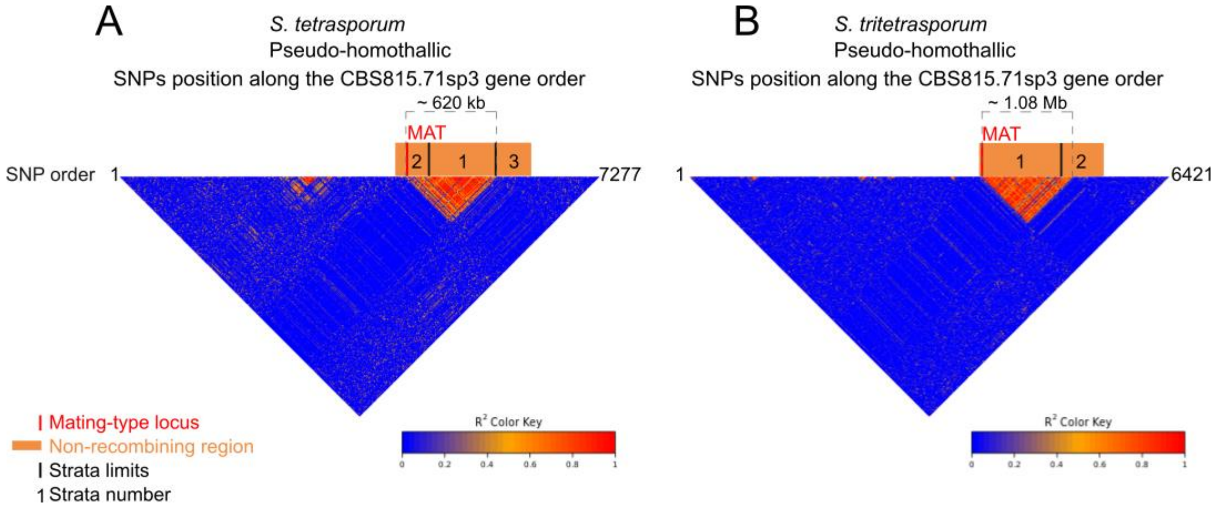

Supplementary Figure 4:

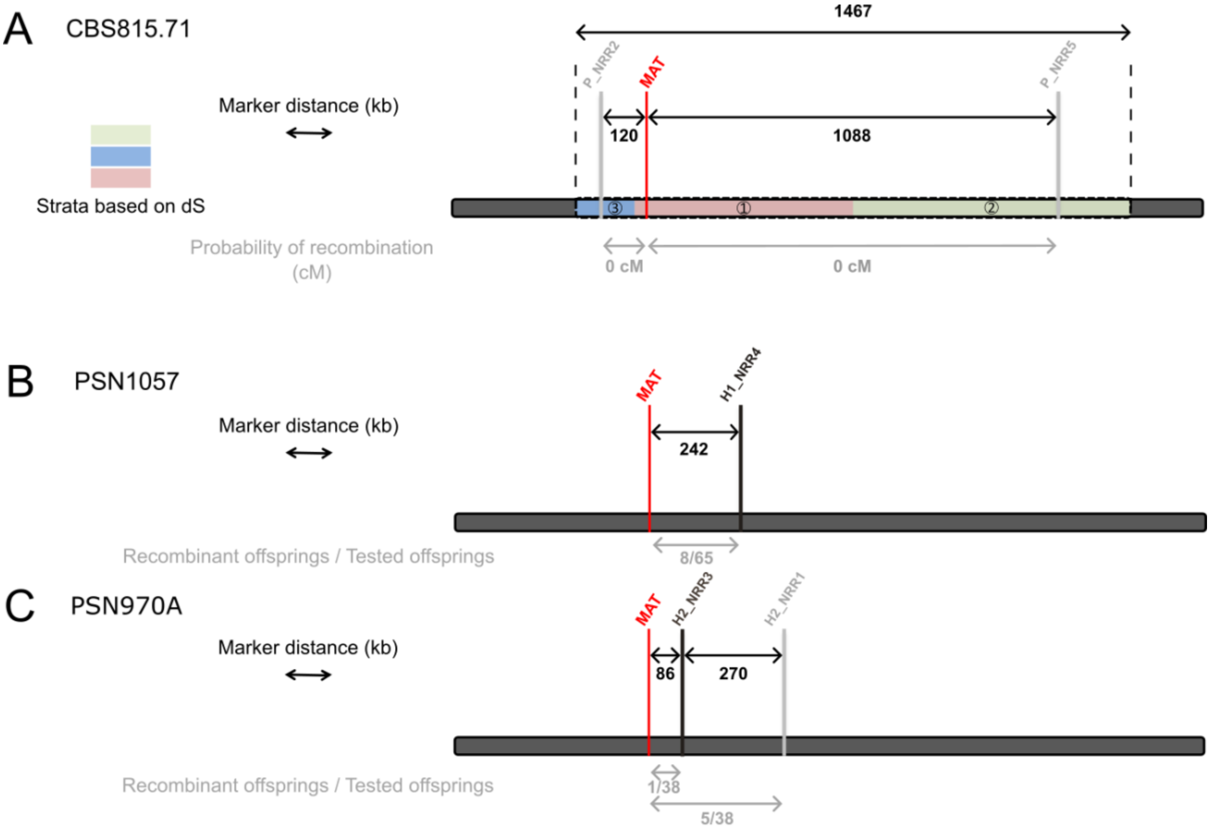
